# Interactions of encoding and decoding problems to understand motor control

**DOI:** 10.1101/811356

**Authors:** Shixian Wen, Allen Yin, Li Zheng, Laurent Itti

## Abstract

Learning a map from movement to neural data (Encoding Problem) and vice versa (Decoding Problem) are crucial to understanding motor control. A principled encoding model that understands underlying neural dynamics can help better solve the decoding problem. Here, we develop a new generative encoding model leveraging deep learning that autonomously captures neural dynamics. After training, the model can synthesize spike trains given any observed kinematics, under the guidance of the learned neural dynamics. When neural data from other sessions or subjects are limited, synthesized spike trains can improve cross-session and cross-subject decoding performance of a Brain Computer Interface decoder. For cross-subject, even with ample data for both subjects, neural dynamics learned from a previous subject can transfer useful knowledge that improves the best achievable decoding performance for the new subject. The approach is general and fully data-driven, and hence could apply to neuroscience encoding and decoding problems beyond motor control.

## Introduction

To understand motor control, one often distinguishes between two crucial problems. Given a goal for complex movement (e.g., specific limb kinematics), the “encoding” problem concerns how to generate neural spike trains to activate muscles and execute the plan. The encoding model needs to learn how to find a good set of neural dynamics, how they interact, and how they may be adjusted and combined for different tasks and behaviors. Conversely, the “decoding” problem seeks to estimate or recover a motor plan from given neural spike trains. Solving the problem of how to decode new and complex movements from neural spike trains may benefit from a formalism that includes neural dynamics learned from the encoding problem.

Much previous research has tried to demystify the encoding and decoding problems. The encoding problem^1–3^ can be formulated as finding appropriate statistical models that assign a conditional probability, *P(D*|*x)*, to any possible neural response *D*, given the desired movement plan *x*. However, previous studies^4–6^ have imposed a strong prior mathematical model onto *P(D*|*x)*, which limits their generality. To tackle the decoding problem, for brain-machine interfaces^7^, one needs to learn the mapping from recorded neural spikes to limb or artificial actuator kinematics. Several methods have been proposed to solve this problem, exploiting Wiener Filters^8,9^, Kalman Filters^10,11^, Particle Filters^12,13^, Point Process methods^14,15^, and state-of-the-art Long Short-Term Memory (LSTM) networks. While powerful, these methods, require a large amount of neural data to achieve good performance^16,17^.

Cross-session and cross-subject scenarios are crucial to designing a commercial brain computer interface (BCI) decoder. In the cross-session scenario, a decoder trained with data from one recording session is used with data from another session with a possibly different number of neurons. This scenario is important for designing a fast adapting BCI decoder, as scarring^18,19^, motion, neural plasticity^20^, or cell death^19^, may change the effective number of recording channels from day to day. In the cross-subject scenario, a decoder is trained with data from one subject and used with data from another subject, usually also with a different number of neurons. Leveraging data from the first subject can be beneficial in some cases, such as obtaining neural activity and covariates of interest from the second subject is challenging, expensive, or impossible^21^. In addition, neural data from the first subject might be inherently easier to decode (e.g., the quality of signal collected by the implanted electrode arrays might be better) than the neural data from the second subject. Even in cases where ample neural data may be available for both subjects, neural dynamics learned from the easier subject might transfer some useful knowledge that improves the decoding performance of the harder subject. However, existing BCI decoders usually fail to generalize to different sessions^22^ or subjects, because they fail to capture the underlying neural dynamics^23^. We believe this is because current approaches lack a principled representation of neural dynamics, obtained through exploration of possible interactions between encoding and decoding problems.

The intuition of this paper is that building a better encoding model that understands neural dynamics can help better solve a decoding problem commonly encountered in brain computer interface experiments. Here we propose a generative model leveraging Deep Learning^24–30^ (Methods) that creates a direct mapping from kinematics to neural data. Contrary to previous approaches, this model does not rely on any strong prior, but instead learns neural dynamics end-to-end from the training data. After learning from kinematics and associated spike trains, we evaluate the learning process by showing that the model is able to synthesize spike trains, given kinematics in the training set (footnote ^1^), that has realistic characteristics (position activity maps, velocity neural tuning curves, and histogram of mean firing rates). In addition, the encoding model can generalize to novel situations, producing synthesized spike trains which we show are sufficient for practical use in a decoder, even though they may not be perfect in all aspects. To show how the encoding model can help better solve the decoding problem, we fine-tune it with small amounts of neural data from another session or subject, thereby quickly adapting^31,32^ it to synthesize spike trains for a different number of neurons, a different day, and possibly a different monkey. In the cross-session scenario, we show that training a BCI decoder on a combination of a limited amount of neural data and the synthesized spike trains can yields higher decoding performance on an independent test set of additional neural data, compared to training the BCI decoder only on the limited neural data. We confirm that this approach also yields better performance than three alternative data augmentation methods (Methods). In the cross-subject scenario, even with ample neural data for both subjects, training a BCI decoder on a combination of the neural data from the second subject and synthesized spike trains derived from the first subject does not impair and sometimes improves beyond the best achievable decoding performance using the neural data from the second subject only. This is because neural dynamics learned from the easier to decode subject can transfer some useful knowledge that may improve the decoding performance for the harder to decode subject. In addition, when the neural data from the second subject is limited, synthesized spike trains that capture the neural dynamics improve the cross-subject decoding performance on some aspects of kinematics. The good performance of the BCI decoder further validates the good quality of the encoding model. For the first time, our results show how one can leverage a deep learning model to effectively enable neural decoding across sessions and subjects.

Recently, Pandarinath et al.^22^ proposed a deep learning generative approach to infer neural population dynamics using sequential auto encoders (LFADS). Our method is a complementary approach to their work. The key difference is that LFADS focuses on creating a mapping from the neural data to low-dimensional latent variables, and then on reconstructing the same neural data from the latent variables. Our method creates a direct mapping from kinematics to neural data. We argue that this direct mapping is an encoding model of how our brain works under the current task. More differences between the two approaches are described in the discussion section.

## Results

### Experimental setup and data preparation

Two monkeys (Monkey C and Monkey M) were chronically implanted with electrode arrays (Blackrock microsystems) in the arm representation of primary motor cortex (M1). We recorded from these electrodes while the monkeys made reaching movements to a sequence of randomly-placed targets appearing on a computer screen^33^. The monkeys were seated in front of the screen and grasped the handle of a planar manipulandum that controlled the position of a cursor. After the cursor reached a given target, a new target appeared, to which the monkeys could reach immediately (Fig. 1a).

**Figure 1:**
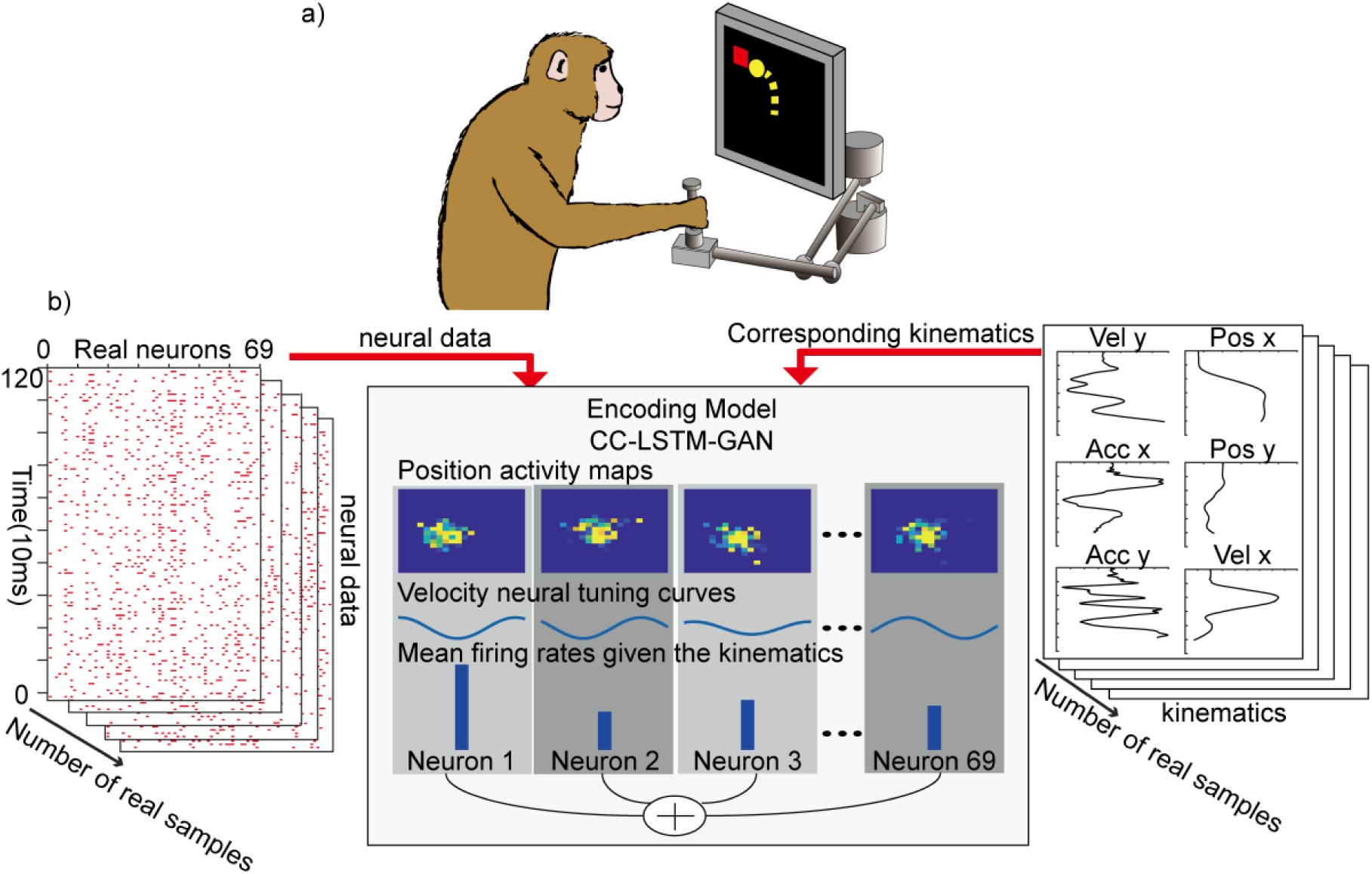
a) Experiment demonstration. b) Train neural encoding model (cc-LSTM-GAN) on neural spikes from Monkey C, letting it learn neural dynamics (position activity maps, velocity neural tuning curves) and synthesize spike trains with a realistic histogram of mean firing rates given the kinematics in the training set.

In the first set of the experiments we analyzed one session of neural data with 33.3 minutes and 69 neurons from Monkey C. We parsed and binned all neural and kinematics data with 10ms time resolution.

### Encoding Problem

To characterize the properties of each neuron in motor cortex, one often collects spike trains from the neurons and calculates properties such as firing rates, position activity maps, and velocity neural tuning curves. We first trained the neural encoding model to synthesize spike trains by presenting two inputs to the model: the kinematics of a movement, and a vector of Gaussian noise which allows the model to generate many different instances of neural spike trains (for different instances of the noise vector). In essence, the neural encoding model has learned a mapping from noise and kinematics to neural spike trains. We can characterize the properties of synthesized spike trains to characterize each *virtual neuron* of the model in the same way as one characterizes real neurons. We found that the neural encoding model learned realistic neural dynamics (position activity maps, velocity neural tuning curves), comparable to real neurons (Fig.2, Fig.3). Given any kinematics in the training set, we confirmed that the neural encoding model synthesized spike trains with a realistic histogram of mean firing rates of virtual neurons compared to real neurons (Fig.3), as further detailed below.

#### Encoding model learned position activity maps

We first asked whether the virtual neurons had position activity maps (activity as a function of position in the workspace) that resembled those of real neurons. This would indicate that the neural encoding model captured how neurons encode position information in the M1 area.

To answer this question, we analyzed neurons’ activity as a function of position. We compared the position activity maps built from synthesized spikes trains and real position activity maps built from neural data. The position activity map is calculated by counting the number of neural spikes across time for different end effector positions and normalizing with respect to the averaged spike counts across positions. Fig. 2a is the normalized real position activity map for real neuron 35. Fig. 2b is the normalized generated position activity map for virtual neuron 35. Fig. 2c is the normalized real position activity map for real neuron 3. The mean square error between the position activity maps Fig. 2a to Fig. 2b is 0.0086 (high similarity) while the mean square error between Fig. 2a to Fig. 2c is 0.4448 (lower similarity). Fig. 2d shows a histogram of mean squared error between real position activity map and generated activity map for each neuron. The mean squared error histogram is right-skewed, and, for 61 out of 69 neurons’ (88.4%), the mean squared error between real position activity map and generated position activity map is less than the average mean squared error between real position activity maps. This shows that, with respect to position activity maps, the model has learned realistic virtual neurons. To show the difference between position activity maps, we plot all maps (Fig. 2e). Note how, in Fig. 2, the position activity maps for neurons in M1 exhibited a strong center bias and limited variability. This is consistent with prior reports that neurons in M1 only weakly represent a conditional probability distribution of the reaches^33^. Dorsal premotor cortex, rather than M1, may exhibit stronger activity when the monkey’s reaches is in a neuron’s preferred direction^33^. Thus, our neural encoding model learned position activity maps that resembled those of real neurons.

**Figure 2:**
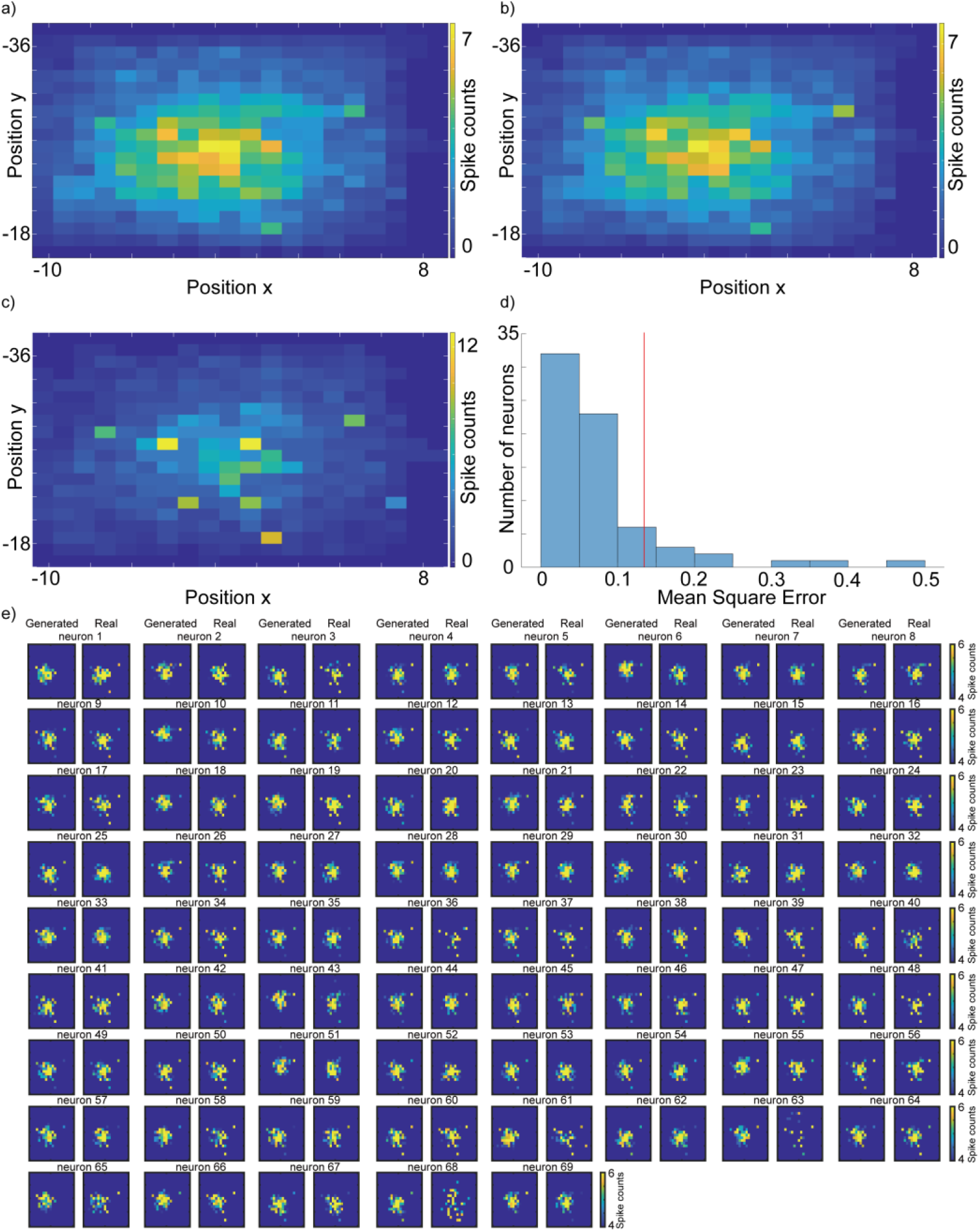
Normalized position activity map, constructed as the histogram of neural activities as a function of position. a) Normalized real position activity map for real neuron 35. b) Normalized generated position activity map for virtual neuron 35. c) Normalized real position activity map for real neuron 3. d) Histogram of mean square error between real position activity map and generated activity map for each neuron. The red line is the averaged mean square error (0.1344) between real neurons. It provides us an upper bound for the difference between real and virtual neurons. The difference from a) to b) is 0.0086. the difference from a to c is 0.4448. The mean squared error histogram is right-skewed, and, for 61 out of 69 neurons’ (88.4%), the mean squared error between real position activity map and generated position activity map is less than the average mean squared error between real position activity maps. e) Position activity maps for all virtual and real neurons with clipped color bar.

#### Learned velocity neural tuning curve

We then asked whether the virtual neurons’ activities had a similar velocity neural tuning curve as the real neurons (Fig. 3a-f). We calculated the hand velocity direction for each 300ms and calculated the spike counts during that 300ms. We plotted the spikes counts vs hand velocity direction for each real and virtual neuron. To calculate the velocity neural tuning curve, we fit a cosine function in velocity space. Fig 3. a-d show that our synthesized spikes trains have similar velocity neural tuning curve shapes for every virtual and real neuron pair. However, from the heatmap, comparing Fig. 3a and Fig. 3b, real neurons exhibit a better systematic structure than virtual neurons (larger amplitude of the tuning curve). The preferred directions of virtual neurons are similar to those of real neurons Fig. 3e, f. The encoding model captures most of the important preferred directions which neurons are tuned to. Thus, our encoding model learned velocity neural tuning curves that resembled those of the real neurons.

**Figure 3:**
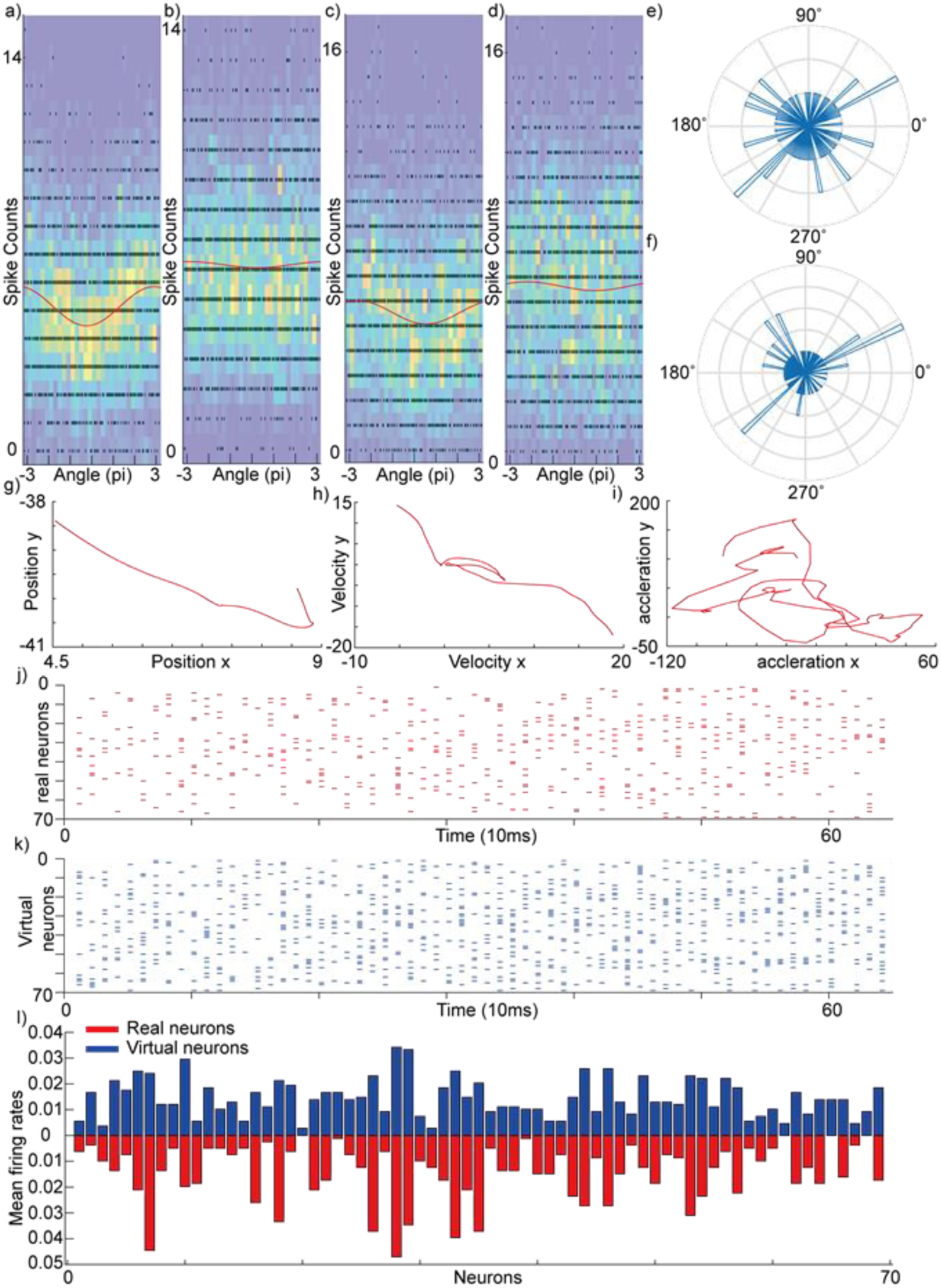
For a-d), we calculated the hand velocity direction for each 300ms and calculated the spike counts during that 300ms. We plotted the spikes counts vs hand velocity direction for each real and virtual neuron. The red line is the velocity neural tuning curve fitted by a cosine function. The black dot is the spike counts for each bin at each angle. The heatmap counts how many black dots are in an area. a) real velocity neural tuning curve for neuron 32. b) generated velocity neural tuning curve for neuron 32. c) real velocity neural tuning curve in velocity space for neuron 57. d) generated velocity neural tuning curve for neuron 57. e) histogram of preferred direction for real neurons. f) histogram of preferred direction for virtual neurons. g-i) position, velocity and acceleration from a trial j) real neuron spikes for this trial k) synthesized spikes trains for this trial l) mean firing rates during this trial for real and virtual neurons.

#### Learned to synthesize spike trains with a realistic histogram of mean firing rates

We asked whether the neural encoding model can learn to synthesize spike trains with a realistic histogram of mean firing rates given a specific kinematic in the training set. We fed the kinematics of a trial in the training set to the encoding model. The encoding model could synthesize neural spikes from kinematics (Fig. 3g, h, i) of the trial. We normalized the summation for each neuron from the synthesized neural spikes (Fig. 3k) and neural data (Fig. 3j). By comparing the mean firing rates for real and virtual neurons over a trial (Fig. 3i), the neural encoding model produces a pattern of firing rates over the population of virtual neurons that is not distinct from the distribution of from real neurons (p >= 0.1019, Kolmogorov–Smirnov test ^34^). Further, we did a bootstrapping test with 1 million samples. 80.43% of samples did not reject the null hypothesis from the Kolmogorov–Smirnov test under 5% significance level. In general, a statistical test cannot conclude that two distributions are identical because there could be minor differences in the distributions, but so small that tests cannot really find the difference^35^. We can only conclude that our sample gives us no evidence against the null hypothesis that the two distributions are the same. Thus, we argue that the patterns of mean firing rates of virtual and real neurons are similar for all practical purposes.

### Decoding problem

In the second experiment, we analyzed an additional session from Monkey C (session 2) with 36.7 minutes and 77 neurons. We also analyzed one session from a new Monkey M – session 1: 11.4 minutes with 60 neurons. We applied the same pre-processing routines as in the previously-analyzed session.

Turning to the decoding problem, previous literature did not consider encoding and decoding models together, with some efforts focusing on encoding and others in decoding. In contrast, we view encoding and decoding models as strongly interacting - a good encoding model that understand neural dynamics can help better solve a decoding problem. Here, we consider two interacting systems: our neural encoding model and a BCI decoder. A good encoding model can synthesize spikes trains from kinematics under the guidance of learned neural dynamics. In the cross-session scenario, when the neural data from another session is limited, we posit that, with the help of synthesized spike trains that captures neural dynamics, we can train a BCI decoder that has a better decoding performance. In the cross-subject scenario, even with ample neural data for both subjects, the neural dynamics learned from one subject (easier to decode) can transfer some useful knowledge to improve the decoding performance of another subject (harder to decode). Training a BCI decoder on a combination of the neural data (from another subject) and the synthesized spike trains, can improve beyond the best achievable decoding performance on acceleration by using the neural data only. In addition, when the neural data from the second subject is limited, synthesized spike trains that capture the neural dynamics improves the cross-subject decoding performance on some aspects of kinematics. Good decoding performance of the BCI decoder further validates the good quality of the encoding model.

The overview of the decoding problem is depicted in Fig. 4 and has 4 steps.

**Figure 4:**
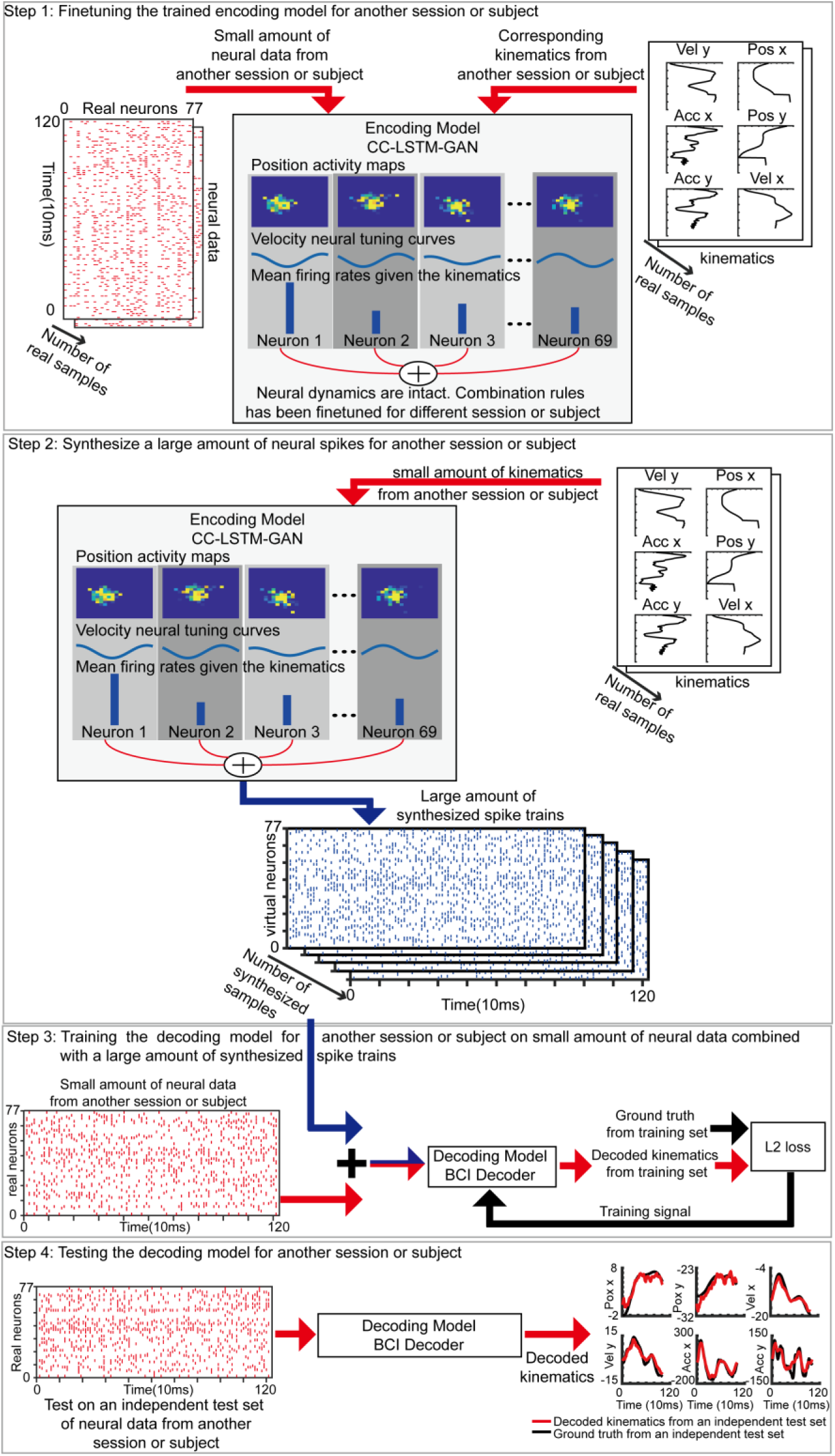
Step 1: Fine-tuning the neural encoding model using small amount of neural data from another session or subject. Step 2: synthesizing a large amount of spike trains using a small amount of real kinematics from another session or subject. Step 3: Combining the large amount of synthesized spike trains with a small amount of neural data from another session or subject to train a BCI decoder. Step 4: Testing the trained BCI decoder on an independent test set from another session or subject.

**Figure 5.**
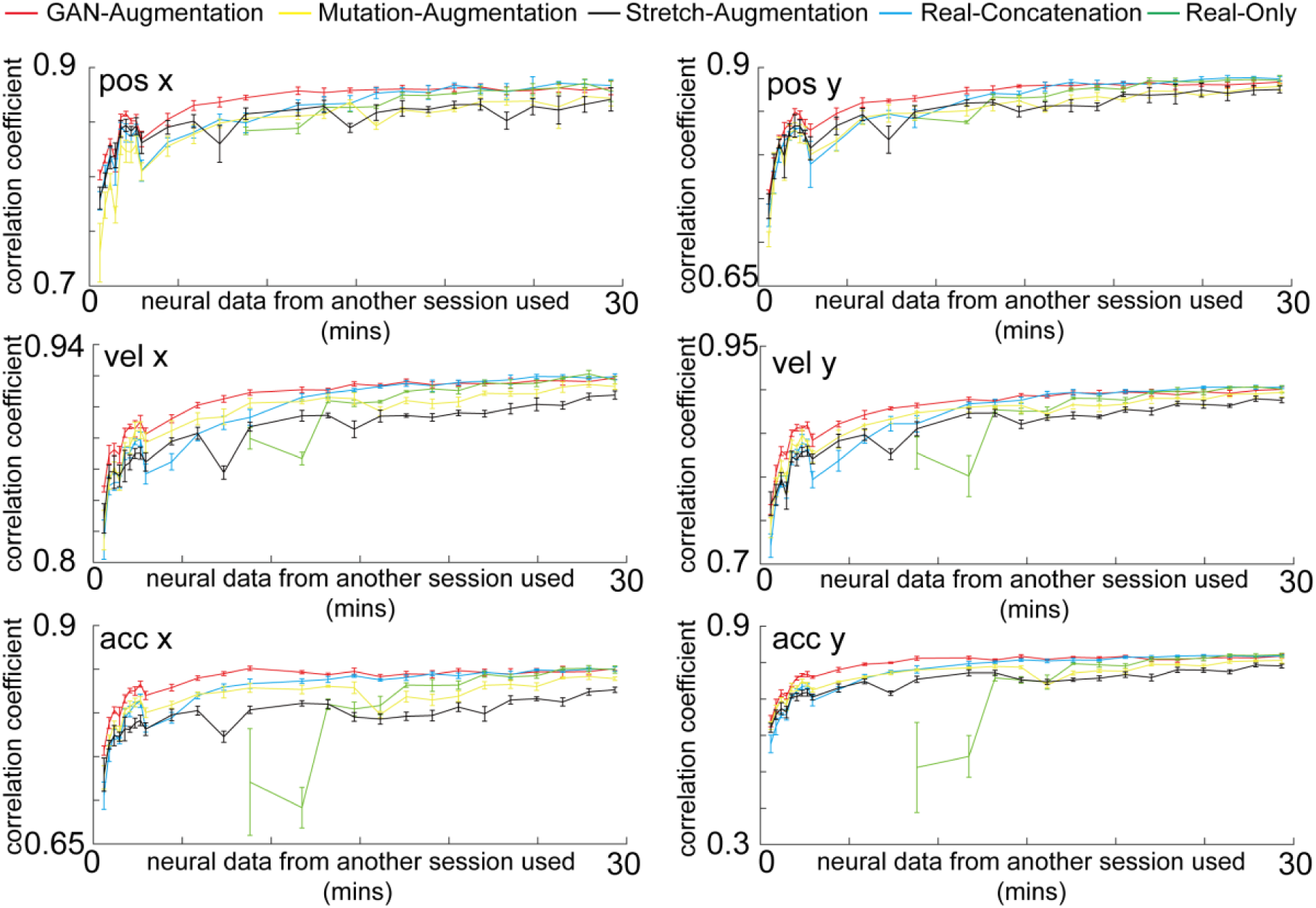
Cross-session decoding. The GAN-Augmentation, Mutation-Augmentation, Stretch-Augmentation, Real-Concatenation and Real-Only methods are shown in red, yellow black, blue and green curves with an error bar. The horizontal axis is the number of minutes of neural data from the session 2 of Monkey C used. The vertical axis is correlation coefficient between the decoded kinematics and real kinematics on an independent test set from the session 2 of Monkey C. Synthesized spike trains that capture the neural dynamics improves the cross-session decoding performance when the neural data from another session is limited.

Step 1: we finetuned (see Methods) the trained neural encoding model using a small amount of neural data from another session or subject. During the finetuning, the neural dynamics are intact. Finetuning only changes the combination rules and number of output neurons. Step 2: we used the finetuned encoding model to synthesize spike trains with real kinematics from another session or subject as inputs. Step 3: we trained a BCI decoder using a small amount of neural data from another session or subject in combination with synthesized spike trains (GAN-Augmentation method). Step 4: we tested the BCI decoder on an independent test set of neural data from another session or subject. We compare the decoding performance from the GAN-Augmentation method with the decoding performance from three other data augmentation methods (Mutation-Augmentation, Stretch-Augmentation, Real-Augmentation, see Methods) and using neural data only (Real-Only, see Methods).

#### Synthesized spike trains improve cross-session decoding performance when training data is limited

With the help of synthesized spike trains, even with very small amount of neural data from another session, we can still achieve a good cross-session decoding performance, which is useful for designing a fast-adapting BCI decoder. In comparison, training a BCI decoder on the same small amount of neural data does not even converge. We trained the neural encoding model on neural data from the session 1 of Monkey C (69 neurons). We used a limited amount of neural data from the session 2 of Monkey C to finetune the neural encoding model, and synthesized spike trains with different number of neurons (77 neurons). From Fig.5, when the neural data from the session 2 of Monkey C is less than 17 minutes, our GAN augmentation method (red curve) is better than the other 3 augmentation methods (blue, yellow, and black curves, p < 0.05) and the Real-Only method (green curve, p < 0.05). if we do not have any augmentation method and train on limited neural data only (Real-Only method), we need at least 8.5 minutes of neural data to make our BCI decoder converge (green). For example, if we only have 35.19 seconds of neural data from the session 2 of Monkey C, the cross-session decoding performance of the GAN-Augmentation method for acceleration x is 0.7569 compared to 0.7053 (7.31% better, p<0.05, Real-Concatenation), 0.7263 (4.22% better, p<0.05, Mutation-Augmentation) and 0.7294 (3.78% better, p<0.05, Stretch-Augmentation) methods. The Real-Only method cannot converge because the neural data from the session 2 of Monkey C is limited. Thus, when the neural data from another session is limited, the neural encoding model can synthesize spike trains to improve the cross-session decoding performance compared to other data-augmentation methods.

#### Transferring learned dynamics and improving the best achievable decoding performance across subject

The neural dynamics learned from Monkey C can transfer some useful knowledge that helps the decoding performance of Monkey M, even with ample neural data for both subjects. Training a BCI decoder on a combination of the neural data (from another subject) and the synthesized spike trains, can improve beyond the best achievable decoding performance on acceleration by using the neural data only. When neural data are ample for both Monkey M and Monkey C, the decoding performance of acceleration of Monkey C (Fig.5, Real-Only method, green curve) is higher than decoding performance of Monkey M (Fig.6, Real-Only method, green curve). This might come from the quality of signals collected by the electrode arrays, or from the fact that neural data from Monkey C is inherently easier to decode than the neural data from Monkey M. Thus, training an encoding model that learns good neural dynamics from Monkey C might transfer some useful knowledge to help the decoding performance of Monkey M on acceleration. For example, from Fig. 6, when all neural data of Monkey M is available (the neural data of Monkey M is ample), the best acceleration y performance is 0.6442 (GAN-Augmentation, 2.28 minutes of neural data used, red curve), compared to 0.4774 (34.9% better, p <0.05, Real-Only method, 9.12 minutes of real neural data used, green curve), 0.6233 (3.35% better, p<0.05, Stretch-Augmentation Method, 9.12 minutes of neural data used, black curve), 0.5909 (9.02% better, p<0.05, Real-Concatenation, 1.824 minutes of neural data used, blue curve) and 0.5127 (25.64% better, p<0.05, Mutant-Augmentation, 4.562 minutes of neural data used, yellow curve). Thus, even with ample neural data for both subjects, the synthesized spike trains learned from Monkey C transferred some useful knowledge to help the cross-subject decoding performance on the acceleration of Monkey M.

**Figure 6.**
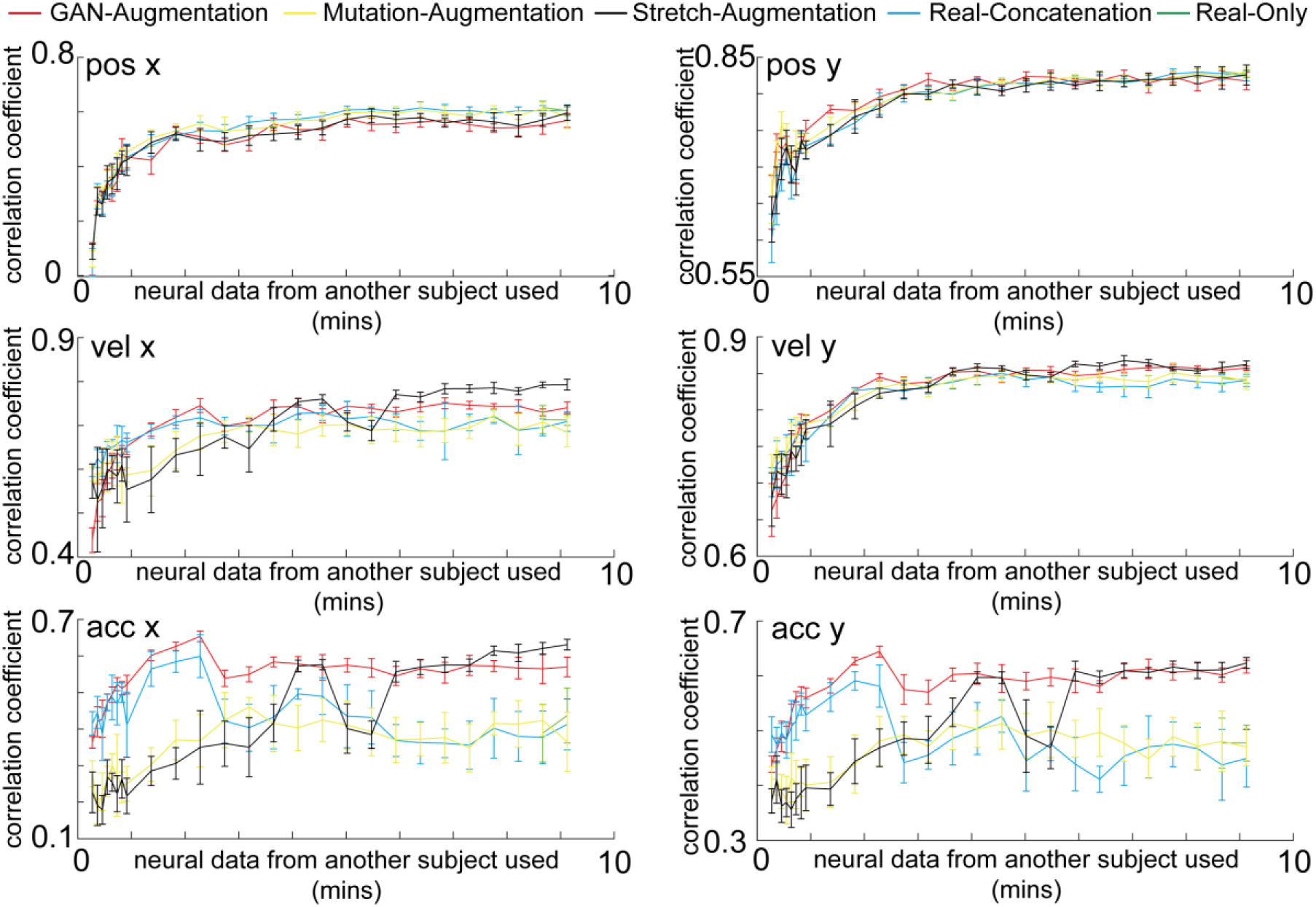
Cross-subject decoding. The GAN-Augmentation, Mutation-Augmentation, Stretch-Augmentation, Real-Concatenation and Real-Only methods are shown in red, yellow black, blue and green curves with an error bar. Cross-subject decoding. The horizontal axis is the number of minutes of neural data from Monkey M used. The vertical axis is the correlation coefficient between the decoded kinematics and real kinematics on an independent test set from the Monkey M. When the neural data from another subject is limited, synthesized spike trains that capture the neural dynamics improves the cross-subject decoding performance on acceleration. Even with ample neural data for both subjects, the neural dynamics learned from one subject can transfer some useful knowledge that improves the best achievable decoding performance on acceleration of another subject.

#### Synthesized spike trains improve cross-subject decoding performance on some aspects of kinematics when training data is limited

In addition, synthesized spike trains that capture the neural dynamics improves the cross-subject decoding performance on acceleration when the neural data from another subject is limited. For example, if we only have 2.28 minutes of neural data from the Monkey M (Fig.6), the cross-subject decoding performance of the GAN-Augmentation method for acceleration y is 0.6442 compared to 0.5812 (10.83% better, p<0.05, Real-Concatenation), 0.4809 (33.95% better, p<0.05, Mutation-Augmentation) and 0.4688 (37.4% better, p<0.05, Stretch-Augmentation) methods.

## Discussion

In the encoding problem, our neural encoding model learned velocity neural tuning curves and position activity maps of virtual neurons that are similar to real neurons. Given any kinematics in the training set, our neural encoding model synthesizes spikes trains that have similar firing pattern as neural data. In addition, the neural encoding model can generalize to novel situations, producing synthesized spike trains which we show are sufficient for practical use in a decoder, even though they may not be perfect in all aspects. In the decoding problem, for the cross-session scenario, with the help of the synthesized spike trains produced by neural encoding model, we could better train the BCI decoder and improve its cross-session decoding performance. For the cross-subject scenario, the synthesized spike trains learned from one subject can transfer some useful knowledge that improves the best achievable decoding performance, on some aspects of the kinematics, of another subject, comparing to training a BCI decoder only on the neural data from another subject. In addition, when the neural data from another subject is limited, synthesized spike trains can improve the cross-subject decoding performance on some aspects of kinematics. Thus, building a better encoding model by understanding underlying neural dynamics can help better solve a decoding problem.

Neural dynamics (position activity maps and velocity neural tuning curves) learned by neural encoding model have a special name in the literature^36–41^ – Motor primitives. Motor cortex is believed to control movement through flexible combinations of motor primitives, elementary building blocks that can be combined and composed to give rise to complex motor behavior. Shadmehr et al. ^40^ defined a motor primitive as the velocity neural tuning curve for each neuron, fitted by a Gaussian function. They built movement trajectories through linear combinations of those velocity tuning curves. In related research, Stround et al.^41^ used gain patterns over neurons or neural groups to predict movement trajectories. Here we use an extended version of Shadmehr’s^40^ definition of motor primitive that includes both position and velocity tuning for each neuron. We hypothesize that our neural encoding model learned motor primitives and their combination rules in an autonomous and principled way. In addition, the encoding model can synthesize corresponding spike trains given kinematics, under the guidance of motor primitives. With a structured representation of motor primitives, and with the consequent help of synthesized spikes trains, we can improve cross-session and cross-subject decoding performance.

In addition, we could interpret the improvement in the decoding performance gained with the encoding model from the perspective of statistics of movements. Kording, et al.^42^ proposed a fundamentally new approach, alignment decoding (DAD), leveraging the statistics of movements. The understanding of the statistics of movements can help us in many situations where obtaining simultaneous recordings of both neural activity and kinematics is challenging, expensive, or impossible. They built prior distributions for feeding, running and reaching tasks. DAD aligned the distribution of its output with statistics of prior distributions to learn a linear decoder. However, to achieve better decoding performance in more complex movements, one would need many prior distributions (high-level templates), because the complex movements might involve sub-part movements such as holding still, rapid reaching, slow reaching, etc. It takes a lot of time to manually craft these templates. It is hard to choose the right form of the templates for a task and to combine them properly. In comparison, our neural encoding model learned neural dynamics such as position activity maps, velocity neural tuning curves (low-level templates) directly from the data. For any complex kinematics, our encoding model could synthesize spike trains (sufficient for practical use) with distributions corresponding to its high-level movements (such as distribution for holding and distribution for rapid reaching) by properly combining those neural dynamics in an autonomous way. There is no need to handcraft high-level templates because they can be constructed from more fundamental low-level templates. Thus, training a BCI decoder to learn those prior distributions contained in the synthesized spike trains can improve the cross-session and cross-subject decoding performance.

Recently, Pandarinath et al.^22^ proposed an interesting method to infer latent dynamics from single-trial neural spike data leveraging an auto-encoder of deep learning (LFADS). Our method is complementary to this work for the following reasons. First, the focus of LFADS is on how to construct the low-dimensional latent variables. Pandarinath et al.^22^ use an auto-encoder that creates a mapping from the neural data to low-dimensional latent variables (neural population dynamics), and reconstructs the same neural data from these low-dimensional latent variables. In comparison, our method uses a generative adversarial network that creates a mapping directly from the kinematics to the neural data. It can synthesize realistic spike trains which demonstrate realistic neural dynamics (position activity maps, velocity neural tuning curve, mean firing rates), given the kinematics in the training set. In addition, it can synthesize novel, but good enough spike trains for practical usage (improve cross-session and cross-subject decoding), given the kinematics from another independent session or subject. Second, LFADS assumes that spikes are samples from a Poisson process. In contrast, our method does not impose any prior distribution on the data and can fit the distribution directly from the data, since a strong prior distribution might limit the generality of the model. Third, a need for stabilization of the latent space arises because of continuous changes in the recording device. Pandarinath et al.^22^ cope with this instability by continuing to train the interface over as long as five months. This may not always be a viable solution in practical applications, because it requires the user to continuously adapt to a changing interface. In comparison, our model only needs one full session (about 24 mins) to achieve stable predictions from kinematics to synthesized spike trains.

Last but not the least, the approach is general and fully data-driven, and hence could be applied to other neuroscience encoding and decoding problems beyond motor control without modifying too many domain specific structures.

## Acknowledgements

This work was supported by the National Science Foundation (grant number CCF-1317433), C-BRIC (one of six centers in JUMP, a Semiconductor Research Corporation (SRC) program sponsored by DARPA), and the Intel Corporation. The authors affirm that the views expressed herein are solely their own, and do not represent the views of the United States government or any agency thereof. We appreciate L.E. Miller and M.G. Perich for providing monkey data.

## Methods

### The BCI decoder

We use the state-of-the-art Long Short-Term Memory (LSTM) network^16,17^ as the decoder. Recurrent neural networks can use their feedback connections to store representation of recent input in the hidden states. However, with the traditional backpropagation through time to update the hidden states, they suffer either gradient exploding or vanishing problem. Long Short-Term Memory creates an uninterrupted gradient flow and thus have a better performance. The structure of LSTM cell can be formalized as

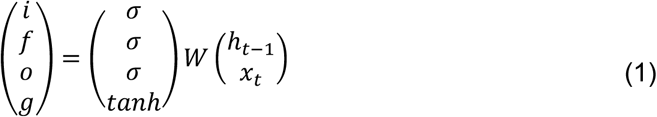

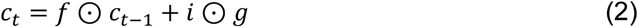

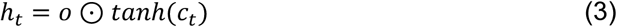

Where *x*_*t*_ is the input at time t. *h*_*t*−1_ is hidden dimension at time t-1. *h*_*t*_ is hidden dimension at time t. *W* is the weight. *σ* is the sigmoid function. i is the input gate, deciding whether to write to cell. f is the forget gate, deciding whether to erase cell. g is the gate gate, deciding how much to write to cell. o is the output gate, deciding how much to reveal cell. *c*_*t*_ is the middle variable. In the LSTM decoder case, we unroll our LSTM cell and consider 200 timesteps for each sample. The input dimension is (N, T, D), where N is the number of samples, T is the number of timesteps, D is the feature dimensions. Our input is batched neural spikes where there are 128 samples, 200 timesteps and number of neurons (69 for session 1, 77 for session 2 of Monkey C, 60 for session 1 of Monkey M) for feature dimensions. The hidden dimensions h is 200 for the LSTM decoder. So, we have an output dimension (N 128, T 200, H 200) from LSTM decoder. We feed this output into a fully connected layer to produce the kinematics (dimension [128, 200, 6]). We apply dropout techniques^43^ and learning rate decay^44^ while training the LSTM decoder.

### Bidirectional LSTM^45^

The output of a sequence at a current time slot not only rely on the sequences before it, but also depends on the sequences after it. So, to better capture the neural dynamics, we use the bidirectional LSTM to build the generator and discriminator in our Constrained Conditional LSTM GAN model. At each time-step t, this network has two hidden state, one for left-to-right propagation and another for the right-to-left propagation. The update rule is

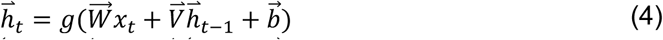

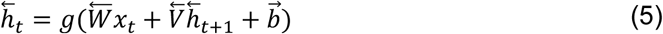

Where 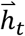 and 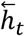 maintains the left-to-right hidden state and right-to-left hidden state separately at time t. g is the LSTM cell update function in Eq.1,2,3.

### Generative adversarial network (GAN)

GANs^25^ provide a tool to learn a map from a random noise input to the desired data distribution in an end to end way, updating its parameters via backpropagation^46^. Thus, it does not require any assumption about the data distribution. It is pure data-driven and does not need a strong prior model which limits the generality. The process of training a GAN can be thought as an adversarial game between a generator and a discriminator. The role of the generator can be thought of as to produce fake currency and use it without detection, while the discriminator learns to detect the counterfeit currency. Competition in this adversarial game can improve both components’ abilities until the counterfeits are indistinguishable from real currency. After this competition, the generator can take the random noise that provides the variations as input and outputs different kinds of realistic bills with different textures. Several approaches have been proposed for image synthesis using GANs enhanced to be able to generate output images for a particular object class, such as conditional GAN^26^, Semi-Supervised GAN^28^, InfoGAN^29^, AC-GAN^27^ and cGANs^30^. In this fake currency scenario, by injecting the conditions (labels of each bill) into the input of GAN, we can select which kind of bills we want to generate (e.g., a 100-dollar bill), the noise provides only the variations of the textures of the bills (e.g. wrinkled, old).

### Constrained Conditional LSTM GAN (cc-LSTM-GAN Supplementary Fig.1)

We propose the constrained conditional LSTM GAN to model the behavior of M1 area given the kinematics. A normal LSTM model takes input which has a dimension of (N, T, D) where N is the number of samples in one minibatch, T is the time horizon, D is the hidden dimension size. We choose 2 seconds as time horizon in our experiments. the first input dimension N is the number of batches. The third input dimension D is the number of neurons for the discriminator or noise dimension for the generator. For each item in the batch, we have a 2 seconds slice of neural spikes with D neurons. Since the number of neurons is the third hidden dimension of LSTM in the discriminator, our discriminator treats different neurons as different individuals that have different neural tuning property. Thus, we call this CC-LSTM GAN encoding model as multiple neural encoding model.

#### Training assistant LSTM decoder (GAN-ta LSTM decoder)

We train a LSTM decoder on neural data from Monkey C beforehand and freeze its parameters when we train our Constrained Conditional Bidirectional LSTM GAN. This decoder applies constrains to the cc-LSTM-GAN. We want to maintain the decoding performance while we train the encoder.

#### Bidirectional LSTM generator

The bidirectional-LSTM generator takes Gaussian noise and real kinematics as input and synthesizes the corresponding spikes trains. We feed the outputs (dimensions [Sample size, Time horizon, Hidden dimension]) of the bidirectional-LSTM into a fully connected layer to synthesize spikes trains with the correct number of neurons (dimensions [Sample size, Time horizon, number of neurons]). We apply 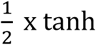 function as the output layer of the fully connected layer which maps a real value to [-0.5, 0.5] that gives us a probability representation of whether the current bin contains a spike event or not. E.g., if the value is 0.3 in the current bin, it means the probability there is a spike event in this bin is 0.8.

#### Bidirectional LSTM discriminator

The Discriminator is a bidirectional-LSTM. It takes the synthesized spikes trains and neural data as input and learns to determine whether a sample is from the neuron data or synthesized spikes trains. We feed the outputs of the bidirectional-LSTM into a fully connected layer to obtain a decision value (0,1). It is a probability that decides whether the current sample is real or fake. In the multiple neural encoding model, we feed the output of a bidirectional-LSTM into another fully connected layer to get the decoded kinematics. This helps us to apply the category constraints.

#### Multiple neural encoding model (CC-LSTM-GAN)

we use a conditional structure that has a GAN category loss to let discriminator tell the difference of both data source distribution (neural spikes distribution) and data labels (kinematics corresponding to this neural spikes) distribution. When the input of bidirectional LSTM discriminator is the neural data (synthesized spikes trains), the real (fake) embedding features are the output of bidirectional LSTM discriminator. GAN embedding category loss is the L2 loss between the real embedding features and the fake embedding features.

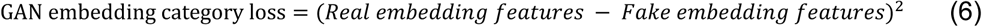

When the input of bidirectional LSTM discriminator is the neural data (synthesized spikes trains), the real (fake) decoded kinematics are the output of the fully connected layer after bidirectional LSTM discriminator. Real (fake) GAN decoding category loss is the L2 loss between real (fake) decoded kinematics and real kinematics. The GAN category loss is the average of real GAN decoding category loss, fake GAN decoding category loss and GAN embedding category loss.

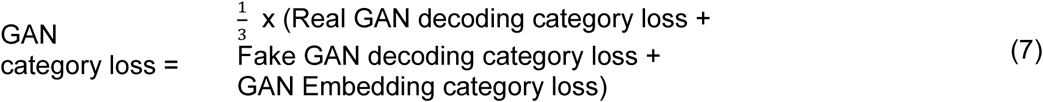

To maintain the source distribution, our cc-LSTM-GAN need to play the min max game, we need to minimize GAN loss discriminator(*L* _*D*_) and GAN loss generator (*L* _*G*_).

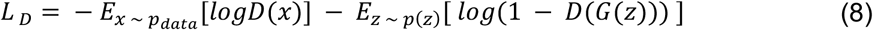

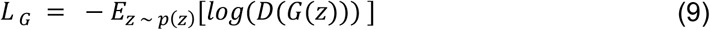

Where *z* is the Gaussian noise, *x* is the neuron spikes, *k* is the kinematics, *p(z)* is the noise distribution, *p*_*data*_ is the data distribution. *L* _*D*_ is the discriminator loss, *L* _*G*_ is the generator loss.

To further maintain the virtual neurons biological structure, we want to maximize the inner product loss between the neural data and synthesized spikes trains. Thus, we have our inner product loss

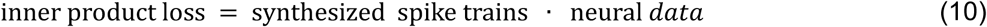

The pre-trained GAN-ta LSTM decoder takes the synthesized spike trains as input and decodes the corresponding decoded generated kinematics. It also takes the neural data as input and decodes the corresponding decoded real kinematics. We apply L2 loss between real kinematics and decoded generated kinematics. We apply L2 loss between decoded generated kinematics and decoded real kinematics. The pre-trained GAN-ta LSTM decoder helps our generator synthesize a more realistic spike trains in terms of the performance of GAN-ta LSTM decoder.

So, the total generator loss is the weighted average of GAN loss discriminator, the L2 loss between decoded generated kinematics and decoded real kinematic, the L2 loss between decoded generated kinematics and real kinematics, and the GAN category loss.

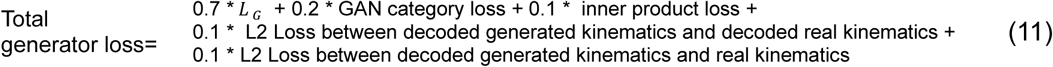

The total discriminator loss is the weighted average of GAN discriminator loss and the GAN category loss. We train this network by real GAN training set and minimize the total discriminator loss and the total generator loss.

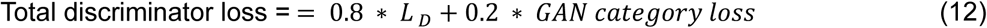

### Finetuning

we train a multiple neural encoding model on the session 1 of Monkey C. In the finetuning process, we take a limited amount of neural data from the session 2 of Monkey C (cross-session) or a limited amount of neural data from Monkey M (cross-subject). We add another fully connected layer on top of the bidirectional LSTM generator and use it to generate the corresponding neural spikes with the same number of neurons as the limited amount of neural data from another session or subject. We freeze the parameter of bidirectional LSTM generator and only trained this new fully connected layer with limited amount of neural data from another session or subject. The loss function of the finetuning process is the inner product loss in Eq.10. Then, we feed the kinematics corresponding this limited amount of neural data into the Generator multiple times to synthesize a large amount of spike trains.

### Data Augmentation

We use multiple data augmentation methods to train a BCI decoder and achieve a better decoding performance than training a BCI decoder on the neural data only.

#### Real-Only method

Without any data augmentation, we train the BCI decoder on the neural data only and test on the neural data of an independent test set. This method requires at least 8.5 mins neural data in the training set to let the BCI decoder converge.

#### Real-Concatenation

We take a limited neural data from the training set and concatenate this limited neural data multiple times until it has the equal or longer length than the whole training set. We train the BCI decoder on the concatenated neural data and test on the neural data of an independent test set.

#### Mutation-Augmentation

We take a limited neural data from the training set. We flip the value of this neural data with 5% probability. We repeat these two steps several times and concatenate the mutated neural data and its kinematics together until it has an equal or longer length than the whole training set. We train the BCI decoder on the mutated neural data and test on the neural data of an independent test set.

#### Stretch-Augmentation

We take a limited neural data from the training set. We stretch the neural data by 10 percent. We fill the empty stretched slots of the neural data by zeros (50% probability) or ones (50% probability). We calculate the average absolute gradients for each kinematics during the transition of each time slot. We fill the empty stretched slots of the kinematics by summation of the positive average absolute gradients (50% probability) or negative absolute gradients (50% probability) and the value of its last slot. We repeat the above steps several times and concatenate the stretched neural data and its kinematics together until it has an equal or longer length than the whole training set. We train the BCI decoder on the stretched neural data and test on the neural data of an independent test set.

#### GAN-Augmentation

we use a limited neural data from the training set for the finetuning process. Then, we combined the synthesized spike trains with the augmented data from Real-Concatenation method. We train the BCI decoder on the combination of synthesized spike trains and concatenated neural data. We test on neural data of an independent test set.

## Supplementary

**Supplementary figure 1:**
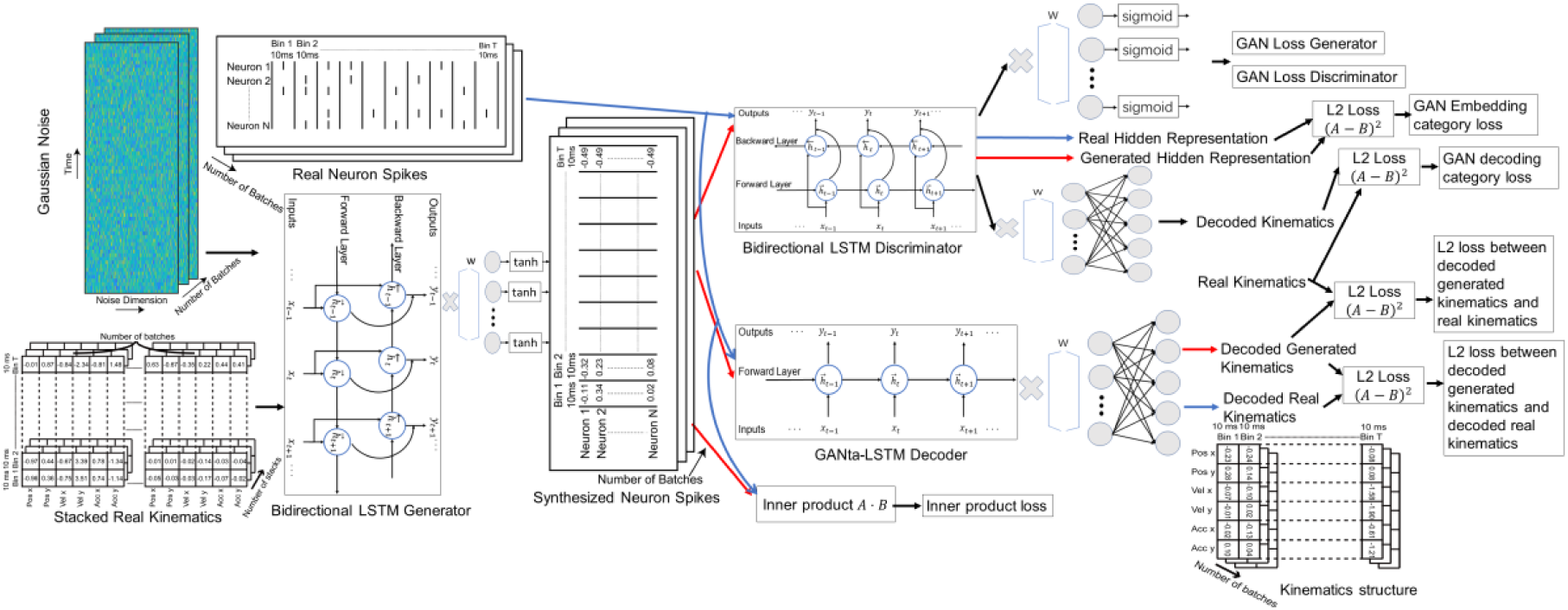
neural encoding model (CC-LSTM-GAN)

## Footnotes

1 Current deep generative models can only generate samples from the distribution they have been trained on. For example, in machine vision, a generative model can only generate images of dogs and cats if it has only been trained on images of cats and dogs. We do not expect it can generalize to images of birds. Here, because each distinct kinematics is a new category such as birds, we do not expect our model to generalize well to novel kinematics in terms of neural dynamics. However, we show that, after fine-tuning with a small amount of neural data from another session or subject, our encoding model can generalize to novel situations improving cross-session and cross-subject decoding performance.

